# The sensitivity of magnetic particle imaging and fluorine-19 magnetic resonance imaging for cell tracking

**DOI:** 10.1101/2021.08.31.458286

**Authors:** Olivia C. Sehl, Paula J. Foster

## Abstract

**Purpose:** Magnetic particle imaging (MPI) and fluorine-19 (^19^F) MRI produce images which allow for quantification of labeled cells. MPI is an emerging instrument for cell tracking, which is expected to have superior sensitivity compared to ^19^F MRI. Our objective is to assess the cellular sensitivity of MPI and ^19^F MRI for detection of mesenchymal stem cells (MSC) and breast cancer cells.

**Methods:** Cells were labeled with ferucarbotran or perfluoropolyether, for imaging on a preclinical MPI system or 3 Tesla clinical MRI, respectively. *In vivo* sensitivity with MPI and ^19^F MRI was evaluated by imaging MSC that were administered by different routes.

**Results:** Using the same imaging time, as few as 4000 MSC (76 ng iron) and 8000 breast cancer cells (74 ng iron) were reliably detected with MPI, and 256,000 MSC (9.01 × 10^16^ ^19^F atoms) were detected with ^19^F MRI, with SNR > 5. *In vivo* imaging revealed reduced sensitivity compared to *ex vivo* cell pellets of the same cell number.

**Conclusion:** MPI has the potential to be more sensitive than ^19^F MRI for cell tracking. We attribute reduced MPI and ^19^F MRI cell detection *in vivo* to the effect of cell dispersion among other factors, which are described.

## Introduction

Cellular imaging with magnetic particle imaging (**MPI**) and magnetic resonance imaging (**MRI**) enables the tracking of cellular therapies and the fate of cancer cells. MPI and Fluorine-19 (**^19^F**) MRI are advantageous as they provide positive image contrast with quantifiable signal, without use of ionizing radiation. This allows for specific and longitudinal tracking of cells and the ability to quantify the number of cells. MRI has been used to image therapeutic cells in patients^1^, whereas MPI is an emerging modality for cell tracking and currently limited to preclinical studies. In this study, we evaluate the sensitivity of preclinical MPI and ^19^F MRI on a clinical system, for imaging of two cell types: mesenchymal stem cells (**MSC**) and breast cancer cells.

Therapeutic mesenchymal stem cells (**MSC**) have great potential for regenerative medicine. There are over 4000 clinical trials ongoing in the U.S, and over 350 in Canada, for stem cell therapies^2^. After administration, many MSC die due to a hostile, pro-inflammatory environment. Importantly, the number of cells that survive and persistence of cells at the implant site provide information on therapeutic status^3^. Critical questions about the safety and success of cell therapies – the delivery, numbers, and persistence of cells – remain unanswered. Cellular imaging has the potential to provide answers, and play a role in optimizing dosage, schedules, and administration routes for cell therapies.

Tracking the fate of cancer cells in preclinical models has been another focus of cellular imaging. Metastasis refers to the spread of cancer from the primary tumor to secondary organs and is the leading cause of death for many types of cancer, including breast cancer. The ability to track the fate of cancer populations provides a powerful tool to study metastasis and potential therapeutics which delay or inhibit metastasis. For cell tracking of MSC and cancer cells, imaging quantification is necessary, and so is the detection of few cells (high cellular sensitivity).

### Sensitivity for MPI

MPI directly detects superparamagnetic iron oxide nanoparticles (SPIONs), which are used as cell labeling agents. Strong magnetic gradients (T/m) are used to localize SPIONs by creating a field-free region (FFR) and oscillating excitation fields (mT) are applied to alter the magnetization of SPIONs present in the FFR. The FFR is traversed across the imaging field of view and the change in SPION magnetization is detected by a receive coil. The resulting image has positive contrast, and the signal is directly related to the amount of SPION and cell number.

The type of SPION and the amount of SPION taken up by cells are two major factors that determine the sensitivity of MPI for cell tracking. Optimal SPIONs for MPI will strongly magnetize with magnetic fields (outside the FFR) and experience fast relaxation rates (at the FFR). Monodisperse, single core SPIONs with core sizes of ~25 nm are considered ideal^4^. Cell labeling by SPIONs is typically conducted through co-culture and is dependent on endocytosis. Carbohydrate coatings, such as carboxydextran, increase the interactions of SPIONs with cell membranes, similarly, transfection agents can be used to coat SPIONs to enhance their incorporation to cells.

The most commonly used SPION for MPI is ferucarbotran (VivoTrax), which is repurposed from the original use as an MRI contrast agent (Resovist). Ferucarbotran has a carboxydextran coat and is a polydisperse agent; some nanoparticles have a core size of 24 nm (30%) and the majority have 5 nm cores (70%). The *in vitro* detection limit using ferucarbotran has been estimated to be approximately 1000 embryonic stem cells (27 pg/cell)^5^. SPIONs designed specifically for MPI are being investigated and show improved performance; *in vivo*, Wang *et al.* (2020) demonstrated detection of 2500 bone mesenchymal stem cells with cubic nanoparticles (29 pg/cell)^6^. Beyond this, detection limits for SPION-labeled cells have not been carefully studied.

### Sensitivity for ^19^F MRI

For ^19^F MRI, cells can be labeled with perfluorocarbon agents such as perfluoropolyether (PFPE) nanoemulsions. Since there is little endogenous ^19^F in biological tissues, these cells can be visualized with high specificity. The signal intensity of these images is directly linear to the number of ^19^F atoms and cell number. The sensitivity of ^19^F MRI cell tracking is impacted by ^19^F cellular loading. PFPE are formulated into nanoemulsions for safe and effective labeling of cells; clinical-grade PFPE agents are available (CS-1000, CelSense Inc.) and have been used in humans^7^. The amount of ^19^F uptake is different for various cell types due to differences in cell size and endocytic ability^8^.

MRI hardware and imaging parameters also play a major role in determining ^19^F sensitivity. Higher magnetic field strengths improve detectability of cells, additionally, the use of specialized coils is integral. Our group has previously demonstrated comparable signal detection at 9.4 T using a birdcage coil compared to 3 T using a surface coil^9^.

^19^F cellular detection limits using various field strengths, hardware, and sequences have been reviewed by Srinivas et al. (2012)^8^. Notably, there were no reported studies at field strengths ≤ 3 T. The translation of cellular MRI techniques to the clinic will require the use of human MRI systems at clinical field strengths. Our group has demonstrated a cellular detection limit of 25,000 PFPE-labeled macrophages *in vitro* at 3T^9^. In the first human clinical trial at 3 T, an *in vivo* cellular detection limit between 1 and 10 million dendritic cells was demonstrated^7^. The sensitivity of ^19^F MRI for detection of MSC and breast cancer cells at 3 T has not been evaluated.

MPI and ^19^F MRI have similar characteristics for cell tracking (positive contrast and quantitation), and MPI is expected to be more sensitive for cell tracking, however, this has not been carefully compared. **The objective of this study**is to assess the *in vitro* and *in vivo* cellular sensitivity of MPI and ^19^F MRI for MSC and breast cancer cells.

## Methods

### Cell Culture

4T1 murine breast cancer cells (Dr. Fred Miller, Wayne State University, MI, USA) were maintained in Dulbecco’s modified Eagle’s medium (**DMEM**) (Gibco, Thermo Fisher Scientific, MA, USA) with 10% fetal bovine serum (**FBS**) and antimycotic/antibiotic. MSCs derived from the bone marrow of C57BL/6 mice (MUBMX-01101 [BE], Cedarlane, Burlington, Ontario, Canada) were cultured in low-glucose DMEM (Thermo Fisher Scientific) with 10% FBS. Cells were maintained at 37°C and 5% CO_2_ and passaged every 2-3 days for 10 days.

### Cell labeling

2 × 10^6^ 4T1 cells or MSCs were seeded for labeling in T75 cm^2^ flasks. After 24 hours, 2.5 mg/mL PFPE nanoemulsion (Cell Sense, Celsense Inc., Pittsburgh, PA, USA) was added to 10 mL complete media and left to co-incubate overnight. Alternatively, cells were labeled with 55 *μ*g Fe/mL ferucarbotran (Vivotrax, Magnetic Insight Inc., Alameda, CA, USA) using transfection agents in a protocol described by Thu *et al.*^10^ Briefly, 60 *μ*L protamine sulfate (stock 10mg/mL) was added to 2.5 mL of serum-free medium, and in a second tube, 20 *μ*L heparin (stock 1000 U/mL) and 90 *μ*L ferucarbotran (stock 5.5 mg/mL) was added to 2.5 mL serum-free medium. These two tubes were individually vortexed, then combined. After adhered 4T1 cells or MSC were washed in PBS, this labeling mix was added to the cells. 4 hours later, 5 mL of complete media was added to cells and left to co-incubate overnight.

### Evaluation of cell labeling

A cytospin of 100 × 10^3^ cells was prepared for all labeled cells which were fixed in 3:1 methanol:acetic acid. Iron-labeled cells were stained with Perl’s Prussian blue (PPB) to identify iron in cells^11^. ^19^F-labeled cells were labeled with nuclear fast red to assess for PFPE nanodroplets^9^. Microscopy of these slides was conducted using the EVOS imaging system (M7000, Thermo Fischer Scientific). The mean intracellular ^19^F content in MSC was measured by NMR using a Varian Inova 400 spectrometer (Varian Inc, Paulo Alto, USA) and methods previously described^9^.

### Preparation of cell pellets for imaging

PFPE-labeled cells or ferucarbotran-labeled cells were washed 3 times in phosphate-buffered saline (PBS) and collected in triplicate samples prepared in a dilution series: 1024, 512, 256, 128, 64, 32, 16, 8, 4, 2, 1, 0.5 (x10^3^) cells. Ferucarbotran-labeled cells were pelleted in 50*μ*L PBS for MPI. ^19^F-labeled cells were pelleted by centrifugation and covered in 50*μ*L agarose. This process was repeated two more times to produce a total of 9 replicates for each cell number and cell type (4T1 or MSC), with each cell labeling agent (PFPE or ferucarbotran).

### MPI of ferucarbotran-labeled cells

All MPI was conducted on a MOMENTUM™ system (Magnetic Insight Inc.) (**Figure 1A**). A small well was secured to the imaging bed, where a PCR tube containing the cell pellet could be placed. This allowed for imaging to occur in the identical location and for the user to easily transfer cell samples. Each cell pellet was imaged individually with the following parameters: field of view (FOV) = 6 cm × 6 cm, gradient strength = 3.0 T/m, dual-channel acquisition (X and Z), excitation amplitude = 22 mT (X-channel) and 26 mT (Z-channel), imaging time = 1.5 minutes. Cell pellets were imaged in descending order of cell number. Once the cell pellet was undetected, 2D imaging with 8 averages (imaging time = 11.8 minutes) and 3D imaging was conducted for all 9 replicates, in attempt to improve MPI sensitivity. 3D images combine 35 projections in a FOV = 6 × 6 × 6 cm (imaging time = 22.8 minutes). If cells were detected in these longer scans, fewer cells were imaged until undetected in 2D (8 averages) and 3D.

**Figure 1.**
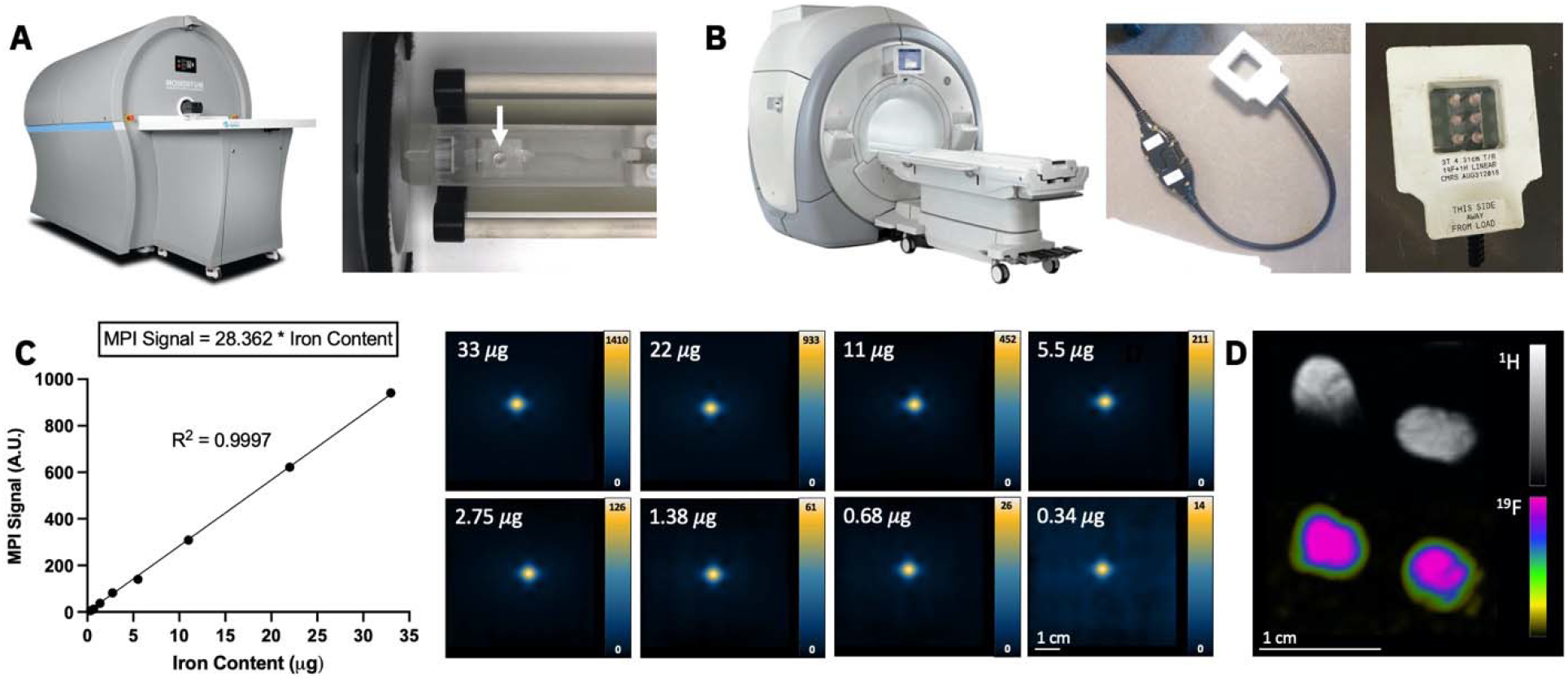
(**A**) Image of a preclinical Momentum MPI and sample holder (white arrow), where individual ferucarbotran-labeled cell samples were placed for imaging. (**B**) Photos of a clinical 3 Tesla MRI with the dual-tuned (^1^H/^19^F) surface coil, used for ^19^F imaging of 6 cell samples. (**C**) A strong linear relationship exists between MPI signal and iron mass (R^2^ = 0.9997). This was determined by imaging multiple samples of ferucarbotran (0.34 *μ*g – 33 *μ*g) and measuring the MPI signal from each sample. The linear equation is subsequently used to convert a measure of MPI signal from ferucarbotran-labeled cells to a measure iron content (*μ*g). (**D**) ^1^H and ^19^F images of two reference phantoms, containing known amount of ^19^F content (3.33 × 10^16^ ^19^F spins/*μ*L), are used for quantification of ^19^F in cell samples.

### ^19^F MRI of perfluorocarbon-labeled cells

^19^F images were acquired on a clinical 3T MRI (Discovery MR750, General Electric) using a 4.31 × 4.31 cm diameter dual tuned (^1^H/^19^F) surface coil (Clinical MR Solutions, Wisconsin) (**Figure 1B**). Six samples of 32, 64, 128, 256, 512, 1024 (x 10^3^) cells (from the same replicate) were imaged at a time using a 3D balanced steady state free precession (bSSFP) pulse sequence. ^19^F imaging parameters were: FOV = 40 × 20 mm, matrix = 40 × 20, slice thickness = 1 mm (1 × 1 × 1 mm^3^ resolution), repetition time/echo time = 5.6 ms/2.8 ms, bandwidth = ±10 kHz, and flip angle = 72°. Flip angle was optimized through a series of ^19^F images (**Figure S1**) and calculated using reported T1 and T2 times at 3T^7,12^. Cell pellets were imaged with 115 excitations (imaging time of 9.5 minutes, or 1.5 minutes/pellet) and 345 excitations (imaging time of 28.3 minutes, or 4.5 minutes/pellet).

### In vivo detection of MSC using MPI and ^19^F MRI

7 NOD/SCID/ILIIrg−/− (NSG) mice were obtained and cared for in accordance with the standards of the Canadian Council on Animal Care, under an approved protocol by the Animal Use Subcommittee of Western University’s Council on Animal Care. In preparation for MPI, 4 mice were fasted for 12 hours to minimize background signal associated with the iron in the mouse digestive system.

To investigate differences in sensitivity *in vivo*, 1 × 10^5^ ferucarbotran-labeled MSC were injected to NSG mice by subcutaneous, intraperitoneal, or intravenous injection. A fourth mouse received 2 × 10^6^ ferucarbotran-labeled MSC by subcutaneous injection. Each mouse was imaged with MPI with a FOV = 12 cm × 6 cm × 6 cm in 2D (2.2 minutes). All other imaging parameters are described above. MPI was conducted immediately before injections, to measure background signal, and immediately after injections. MPI signal for each mouse was calculated as the difference between pre-injection signal (background) and post-injection signal. MPI signal from cells *in vivo* was compared to MPI signal from cell pellets.

Similarly, 2 × 10^6^ PFPE-labeled MSC were injected to NSG mice by subcutaneous or intraperitoneal injection. A third mouse received fewer PFPE-labeled cells (1 × 10^5^) by subcutaneous administration. Following cell administration, ^19^F MRI was conducted for each mouse by placing the surface coil directly above the injection site to maximize sensitivity (shown in **Figure 7E**); the mouse receiving cells by subcutaneous was prone for imaging and the mouse receiving cells by intraperitoneal injection was supine. Imaging parameters are the same as listed above, however with a FOV = 60 × 30 mm with matrix size = 60 × 30 (1 × 1 × 1 mm^3^ resolution), and 200 excitations (imaging time = 35.5 minutes). *In vivo* ^19^F signal was quantified (described below) and compared to signal detected from cell pellets, which were included alongside the mouse.

### Image analysis

For MPI signal calibration, an additional 8 samples of ferucarbotran (5.5 mg/mL) were imaged with identical parameters in a dilution series in PBS: 33, 22, 11, 5.5, 2.25, 1.38, 0.68, 0.34 *μ*g ferucarbotran. A linear relationship was found between iron mass and MPI signal (**Figure 1C**) and the equation of the line was used to calculate associated iron content from each cell sample.

2D MPI of the empty sample holder was conducted at the beginning (0h), middle (3h), and end (6h) of six imaging sessions. The standard deviation of background noise (**SD_noise_**) was measured in these 18 images. To quantify signal from ferucarbotran-labeled cells in pellets and *in vivo*, a threshold of 5 times the average background SD_noise_ was used to mask lower amplitude signal and yield a reliable measurement of MPI signal. This imaging criteria is based on MPI signal with SNR > 5 (Rose Criterion)^13^. Total MPI signal was calculated as mean MPI signal * volume of ROI (mm^2^ or mm^3^).

Delineation of ^19^F signal was conducted using a similar method as MPI (5*SD_noise_). Background SD_noise_ of ^19^F signal for each 3D image was measured by drawing an ROI in background noise of one image slice. Due to Rician distribution observed in background signal noise, ^19^F signal between 5*SD_noise_ and 8*SD_noise_ was corrected using a factor of 0.655, as described by Bouchlaka et al. (2016)^14^ (**Figure S2**). Total ^19^F signal was calculated as mean ^19^F signal * volume of ROI. Two reference phantoms containing 3.33 × 10^16^ ^19^F spins/*μ*L were included in the imaging field of view for calibration (**Figure 1D**). ^19^F content from cell pellets was calculated as:

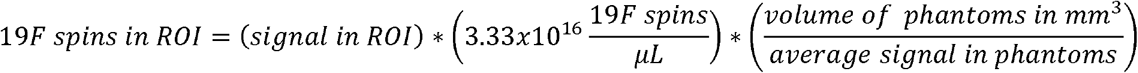

We defined MPI and ^19^F MRI *in vitro* cell detection limits as the minimum number of MSC and 4T1 cells detected with SNR >5. Thus, cells with signal below the 5*SD_noise_ criteria were considered undetected. The amount of ferucarbotran or PFPE associated with the lowest cell number was calculated as iron mass per cell * number of cells or ^19^F atoms per cell * number of cells, respectively.

### Statistical analysis

All statistical analysis were performed using GraphPad Prism version 9. Linear regression was performed for MPI calibration (known iron mass vs. measured MPI signal) to determine the calibration equation. This line is forced through the origin, under the assumption that background MPI signal, without a sample of iron, has an average of 0. Pearson’s correlation was conducted for MPI (number of cells vs. measured MPI signal) and ^19^F MRI (number of cells vs. measured ^19^F signal). Analysis of co-variance (**ANCOVA**) was used to evaluate whether the MPI sensitivity (slope) was significantly different for ferucarbotran-labeled 4T1 cells and MSC (number of cells vs. measured MPI signal). Analysis of variance (**ANOVA**) was used to analyse differences between MPI signal measured from each cell number and again for ^19^F signal measured from each cell number. A p-value of .05 was used to determine statistical significance, unless otherwise indicated.

## Results

### MPI Cellular Sensitivity

The detection of ferucarbotran-labeled cells using 2D MPI (imaging time = 1.5 minutes) is shown in **Figure 2A** (4T1 cells) and **Figure 3A** (MSC). MPI signal measured from samples of 8000 – 1,024,000 4T1 cells and 4000 – 1,024,000 MSC had SNR > 5. The iron mass significantly increased with cell number for both cell types (**Figure 2A, 3A**). Importantly, the standard deviation of background noise, measured three times over six imaging sessions, showed no significant differences (**Figure S3**).

**Figure 2.**
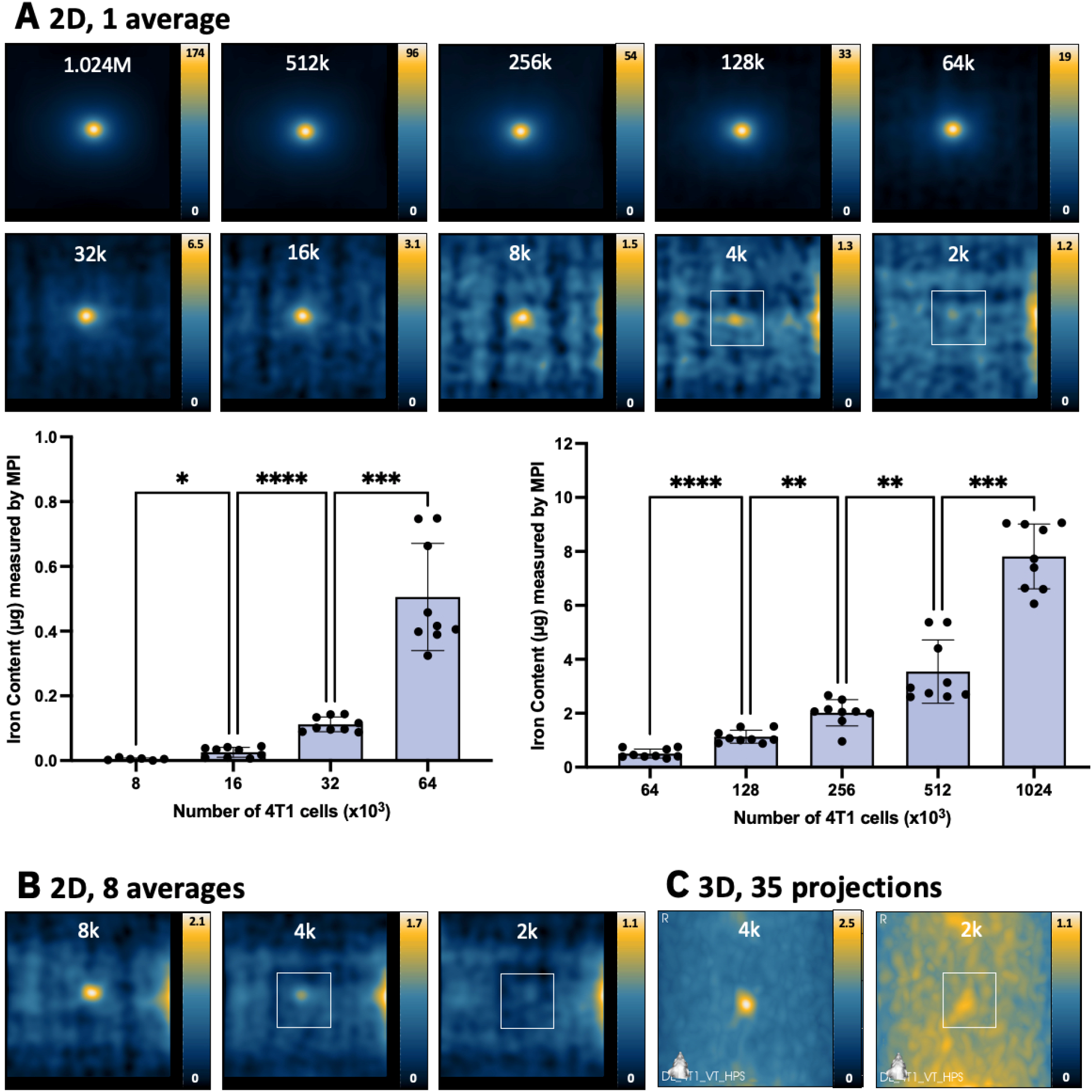
MPI detection of ferucarbotran-labeled 4T1 cells. (**A**) 2D MPI of individual 4T1 cell pellets containing the indicated cell numbers (M = 10^6^, k = 10^3^). As few as 8000 cells (0.074 *μ*g ferucarbotran) could be detected with SNR > 5 in 1.5 minutes. 2D MPI signal (and the associated iron content) significantly increases with the number of 4T1 cells (* p < .05, ** p < .01, *** p < .001, **** p < .0001). (**B**) With 8 signal averages, the same result was found. (**C**) In 3D, the detection of 4000 cells (0.037 *μ*g ferucarbotran) with SNR > 5 was possible in 23 minutes. Images with SNR < 5 have white boxes to indicate the placement of the cell sample.

**Figure 3.**
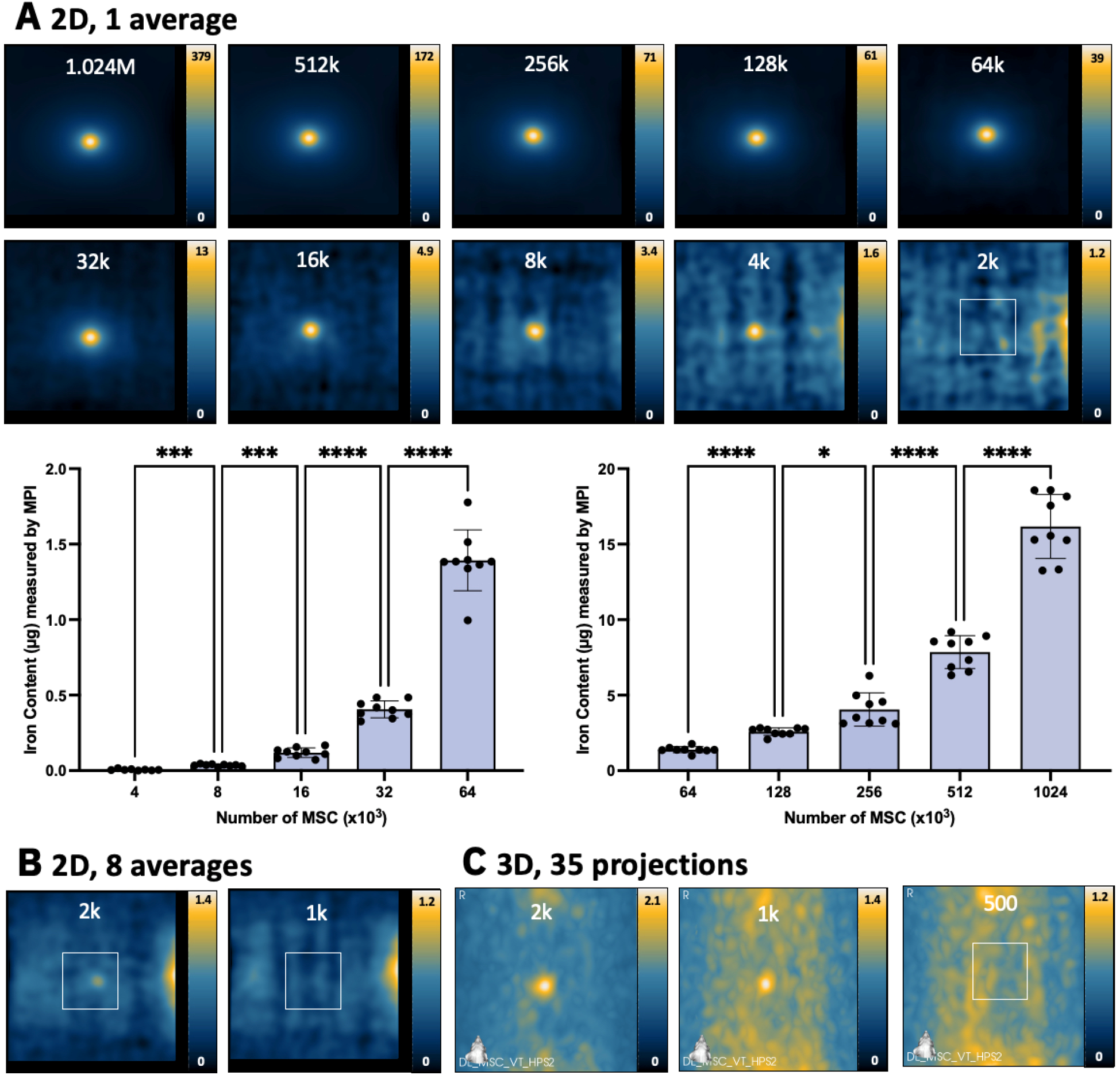
MPI detection of ferucarbotran-labeled MSC. (**A**) 2D MPI of individual MSC pellets containing the indicated cell numbers (M = 10^6^, k = 10^3^). As few as 4000 MSC (0.076 *μ*g ferucarbotran) could be detected with SNR > 5 in 1.5 minutes. 2D MPI signal (and the associated iron content) significantly increases with the number of MSC (* p < .05, ** p < .01, *** p < .001, **** p < .0001). (**B**) With 8 signal averages, the same result was found. (**C**) In 3D, the detection of 2000 cells (0.038 *μ*g ferucarbotran, with SNR > 5) and 1000 cells (0.019 *μ*g ferucarbotran, with SNR > 3) was possible in 23 minutes. Images with SNR < 5 have white boxes to indicate the placement of the cell sample.

MPI signal (and the associated iron content) was strongly correlated with cell number for ferucarbotran-labeled 4T1 cells and MSC (**Figure 5A**). The slope of the line was significantly steeper (factor of 2.07) for MSC compared to 4T1 cells (p < .0001). Enhanced ferucarbotran labeling was measured in MSC (19.09 ± 2.50 pg Fe/cell) compared to 4T1 cells (9.22 ± 1.42 pg Fe/cell), which can be visualized in PPB-stained cells (**Figure 5B**).

With 2D MPI, the lowest cell numbers detected with SNR > 5 were 8000 4T1 cells (6/9 replicates, corresponding to 74 ng iron) and 4000 MSC (8/9 replicates, corresponding to 76 ng iron). All other replicates of 8000 4T1 cells and 4000 MSC had SNR > 3. For both cell types, 2D imaging with 8 averages did not improve cell detection (**Figure 2B, 3B**). However, additional averaging does reduce the standard deviation of background signal from 0.235 (1 average) to 0.183 (8 averages) (p < .0001).

In 3D images (22.8 minutes), detection of 4000 4T1 cells (37 ng iron) was enabled (3 of 9 replicates SNR > 5, and 8 of 9 had SNR > 3) (**Figure 2C**). Improvements to detection of MSC was also seen in 3D; 2000 MSC (38 ng iron) were detected (6 of 9 replicates SNR > 5, and 9 of 9 had SNR > 3) (**Figure 3C**). In 4 of 9 replicates, the detection of 1000 MSC (19 ng iron) was possible in 3D with a threshold of SNR > 3.

### ^19^F MRI Cellular Sensitivity

MSC were labeled with 3.52 ± 1.55 × 10^11^ ^19^F/cell (NMR) and the nanodroplets associated with the ^19^F agent were identified with microscopy (**Figure 5D**). The average number of ^19^F spins determined by NMR was not significantly different from that measured by MRI in both short (p = .3648) and long scans (p = .8541). 4T1 cells did not label sufficiently with PFPE, as ^19^F content in 1 × 10^6^ cells was undetected by NMR. Thus, ^19^F imaging experiments with PFPE-labeled 4T1 cells did not continue.

In ^19^F images (with imaging time = 1.5 minutes/pellet), the range of 256 – 1024 × 10^3^ PFPE-labeled MSC were detected (**Figure 4A**) and ^19^F signal was strongly correlated with cell number (R^2^ = 0.9983) (**Figure 5C**). The average number of cells detected per voxel from these scans was 1.30 ± 0.51 × 10^5^ cells/mm^3^. The lowest cell number detected from these ^19^F scans was 256 × 10^3^ MSC (9.01 × 10^16^ ^19^F atoms, 30.21 mM), which was detected in 4 of 9 replicates with SNR > 5. ^19^F signal measured from 1.024 × 10^6^ MSC was significantly higher than signal measured from 512 × 10^3^ cells (p < .01), which was higher than signal measured from 256 × 10^3^ cells (p < .05) (**Figure 4C**).

**Figure 4.**
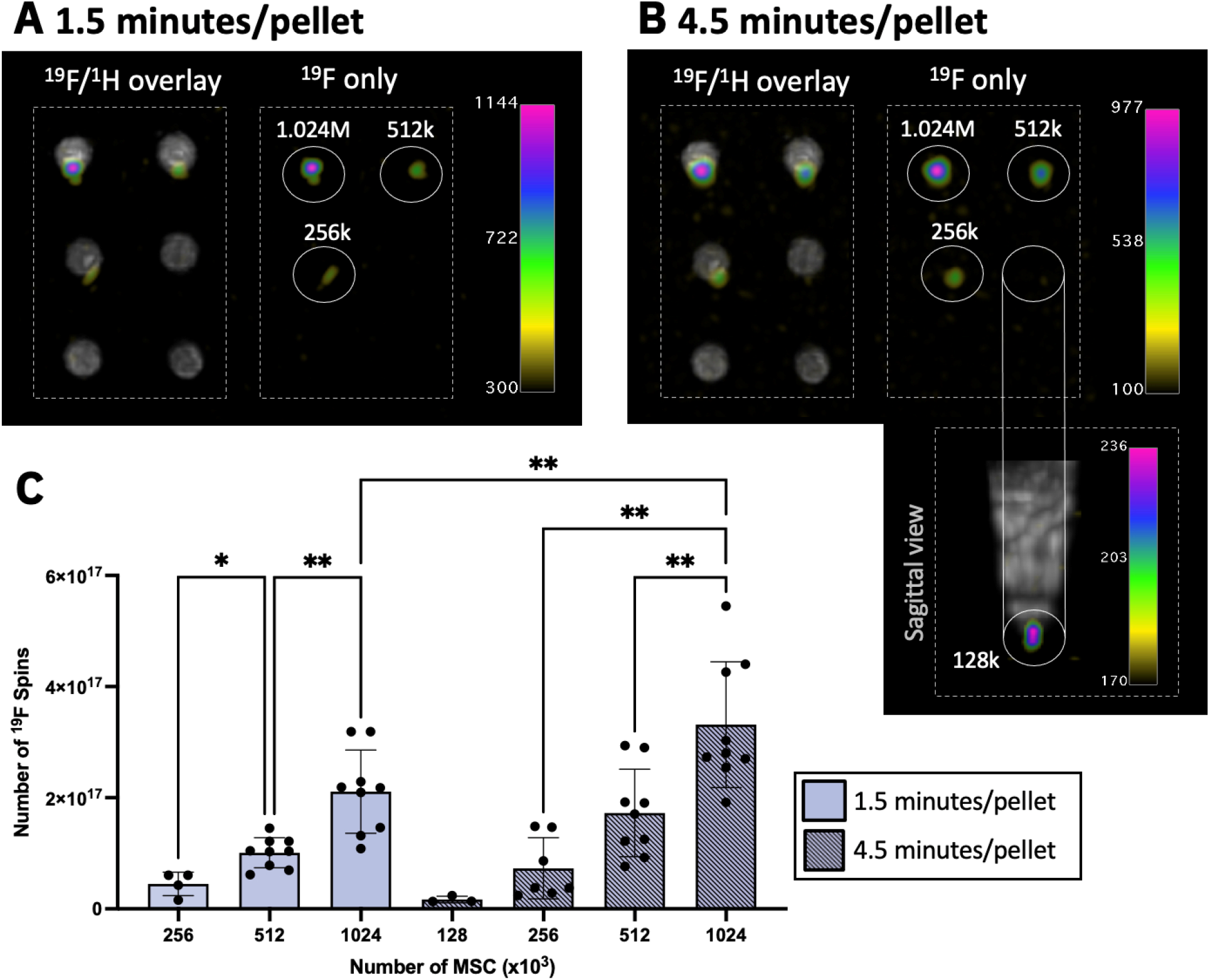
^19^F MRI detection of PFPE-labeled MSC. (**A**) ^19^F images of six samples with various cell numbers (M = 10^6^, k = 10^3^) imaged 1.5 minutes/pellet. As few as 256 × 10^3^ cells (9.01 × 10^16^ ^19^F atoms) could be detected with SNR > 5. (**B**) With longer imaging time (4.5 minutes/pellet), the detection of 128 × 10^3^ cells (4.51 × 10^16^ ^19^F atoms) was possible with SNR > 5. (**C**) Quantification revealed significant differences in ^19^F signal between different numbers of MSC (* p < .05, ** p < .01). Significantly more ^19^F signal was detected from 1.024 × 10^6^ cell samples with longer imaging times (p < .01).

**Figure 5.**
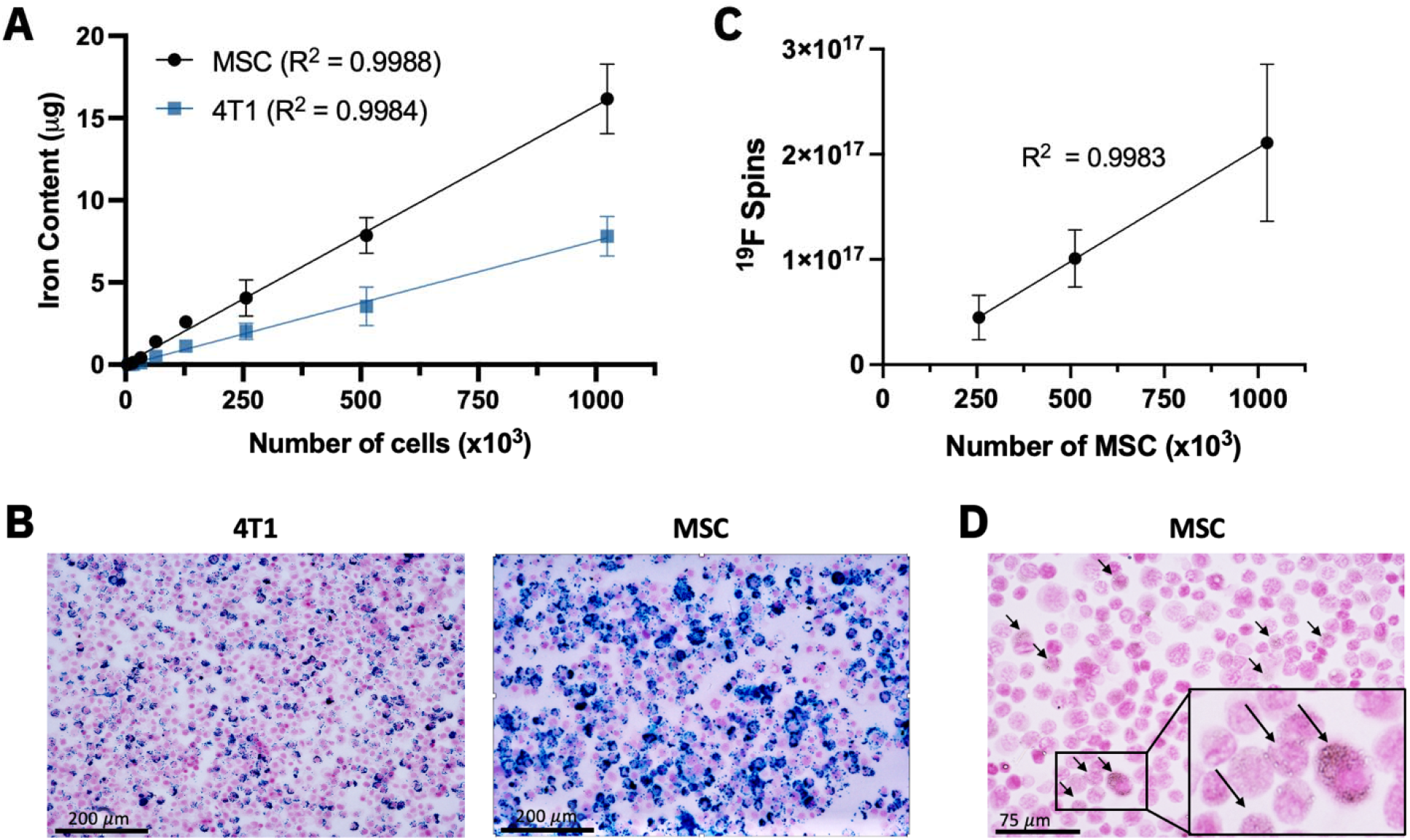
(**A**) The number of ferucarbotran-labeled MSC and 4T1 cells is strongly correlated with iron content measured by MPI (R^2^ > 0.998). The slope of the line for MSC is higher than 4T1 (p < .0001), indicating higher sensitivity and enhanced uptake of iron in MSC compared to 4T1 cells as shown with PPB stain (**B**). (**C**) A strong linear correlation exists between the number of PFPE-labeled MSC and detected ^19^F signal (R^2^ = 0.9983). (**D**) PFPE labeling identified as nanodroplets in microscopy (black arrows).

With longer imaging times (4.5 minutes/pellet), ^19^F sensitivity was improved (**Figure 4B**) and the SD_noise_ was significantly reduced compared to 1.5 minute/pellet scans (p < .0001). The average number of cells detected per voxel from these scans was 8.55 ± 2.97 × 10^4^ cells/mm^3^. As few as 128 × 10^3^ MSC (4.51 × 10^16^ ^19^F atoms, 19.01 mM) were detected with SNR > 5 in 3 of 9 replicates. Additionally, ^19^F signal with SNR > 5 was detected in 7 of 9 replicates of 256 × 10^3^ MSC, which corresponds to 9.01 × 10^16^ ^19^F atoms (28.01 mM). Significant differences in ^19^F signal were measured from 256, 512, and 1024 (x 10^3^) MSC (**Figure 4C**). Furthermore, significantly more ^19^F signal was measured from 1.024 × 10^6^ cells when using longer imaging times (4.5 minutes/pellet) compared to 1.5 minutes/pellet (p < .01).

### *In vivo* sensitivity of MPI and ^19^F MRI

A comparison between MPI signal from a cell pellet and cells *in vivo* was conducted with different injection routes (**Figure 6A**). These MSC were labeled with 28.9 ± 3.4 pg iron/cell. Compared to a pellet of 2 × 10^6^ ferucarbotran-labeled MSC, MPI signal was only reduced by 5% with subcutaneous injection of these cells (**Figure 6 B,C**). Quantification revealed the iron mass measured from the cell pellet (52.98 *μ*g) was similar to what was measured after subcutaneous injection (50.21 *μ*g). However, for 1 × 10^5^ MSC, *in vivo* MPI showed a reduction in MPI signal measured from MSC injected subcutaneously (49%), intraperitoneal (53%), and intravenous (15%), compared to signal from a pellet of 1 × 10^5^ MSC (**Figure 6 D,E**). For 1 × 10^5^ MSC, the measured iron content was 3.13 *μ*g in the cell pellet, compared to 1.52 *μ*g after subcutaneous injection, 1.66 *μ*g after intraperitoneal injection, and 0.48 *μ*g after intravenous injection. Therefore, the iron content measured from 1 × 10^5^ cells *in vivo* was reduced compared to cells in the pellet, despite being the same number of cells. After mouse fasting, the background *in vivo* MPI signal from the mouse digestive system was 25.8 ± 10.0 arbitrary units (**A.U.**) (shown **Figure 6F**). This background signal was accounted for in each mouse by signal subtraction, prior to calculation of MPI signal and iron mass measured from cells.

**Figure 6.**
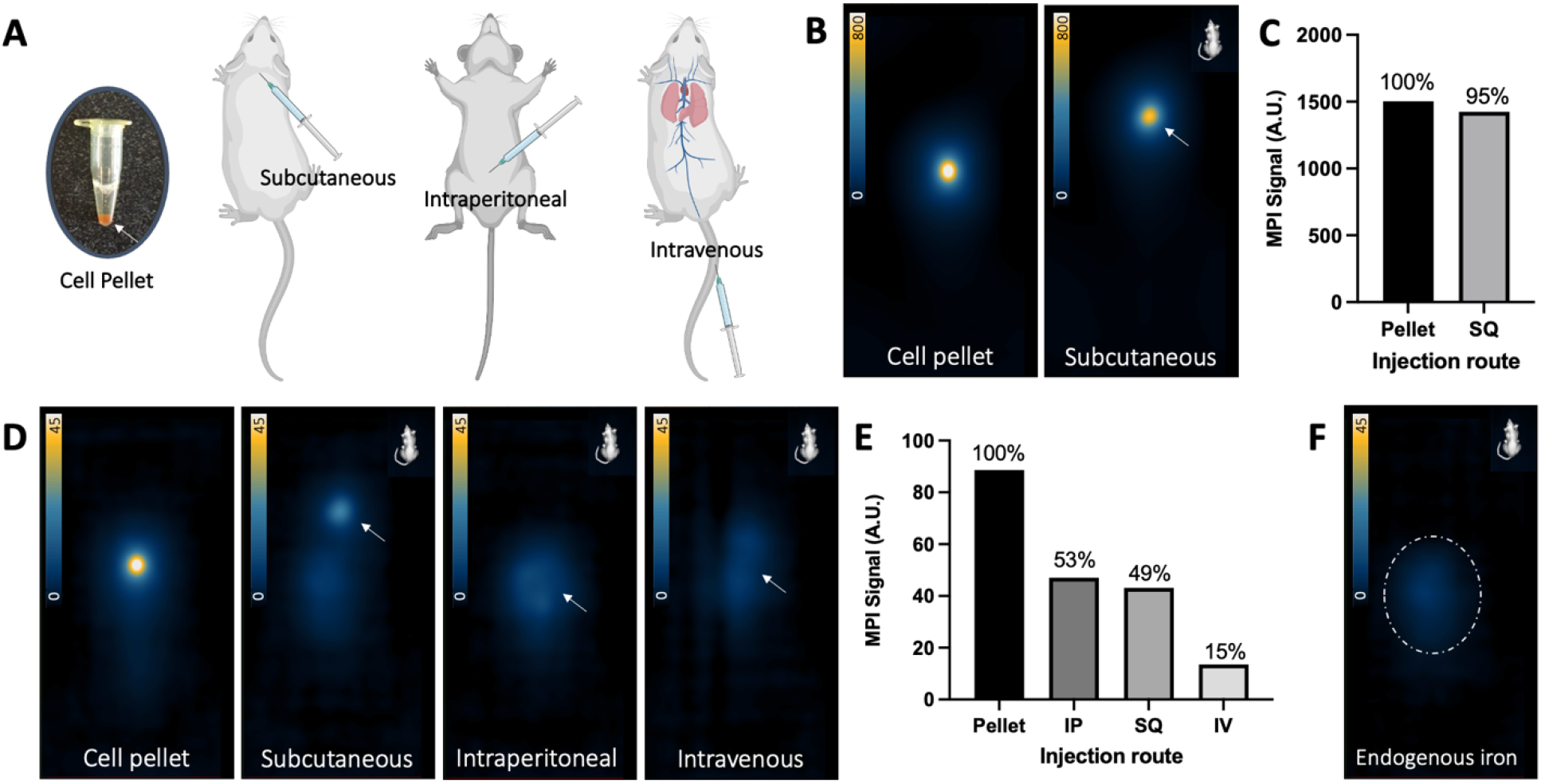
(**A**) Cartoon showing subcutaneous (SQ), intraperitoneal (IP), and intravenous (IV) injection in mice. (**B**) Comparison of MPI signal from 2D scans of 2 × 10^6^ ferucarbotran-labeled MSC in a cell pellet to subcutaneous injection. (**C**) Measured MPI signal from 2 × 10^6^ MSC wa similar *in vivo* and *ex vivo*. (**D**) Detection in 2D of 1 × 10^5^ ferucarbotran-labeled MSC as a cell pellet and after subcutaneous, intraperitoneal, and intravenous injection. (**E**) Measured MPI signal from 1 × 10^5^ cells is reduced *in vivo* compared to signal in the pellet. (**F**) Some background MPI signal exists owing to iron in mouse digestion. This signal was minimized with 12-hour fasting and accounted for by subtracting pre-injection signal from post-injection signal for each mouse.

The detection of 2 × 10^6^ ^19^F-labeled MSC *in vivo* was compared to MSC in a pellet (**Figure 7**). Reduced ^19^F signal (72%) was detected from MSC following subcutaneous injection (**Figure 7A, D**). After intraperitoneal injection, the same number of cells were dispersed and appeared as lower intensity ^19^F signal (**Figure 7B**), however higher ^19^F signal was measured from these cells compared to the pellet by 6.65-fold (**Figure 7D**). 1 × 10^5^ MSC administered subcutaneously were undetected as this cell number is below the detection limit (**Figure 7C**).

**Figure 7.**
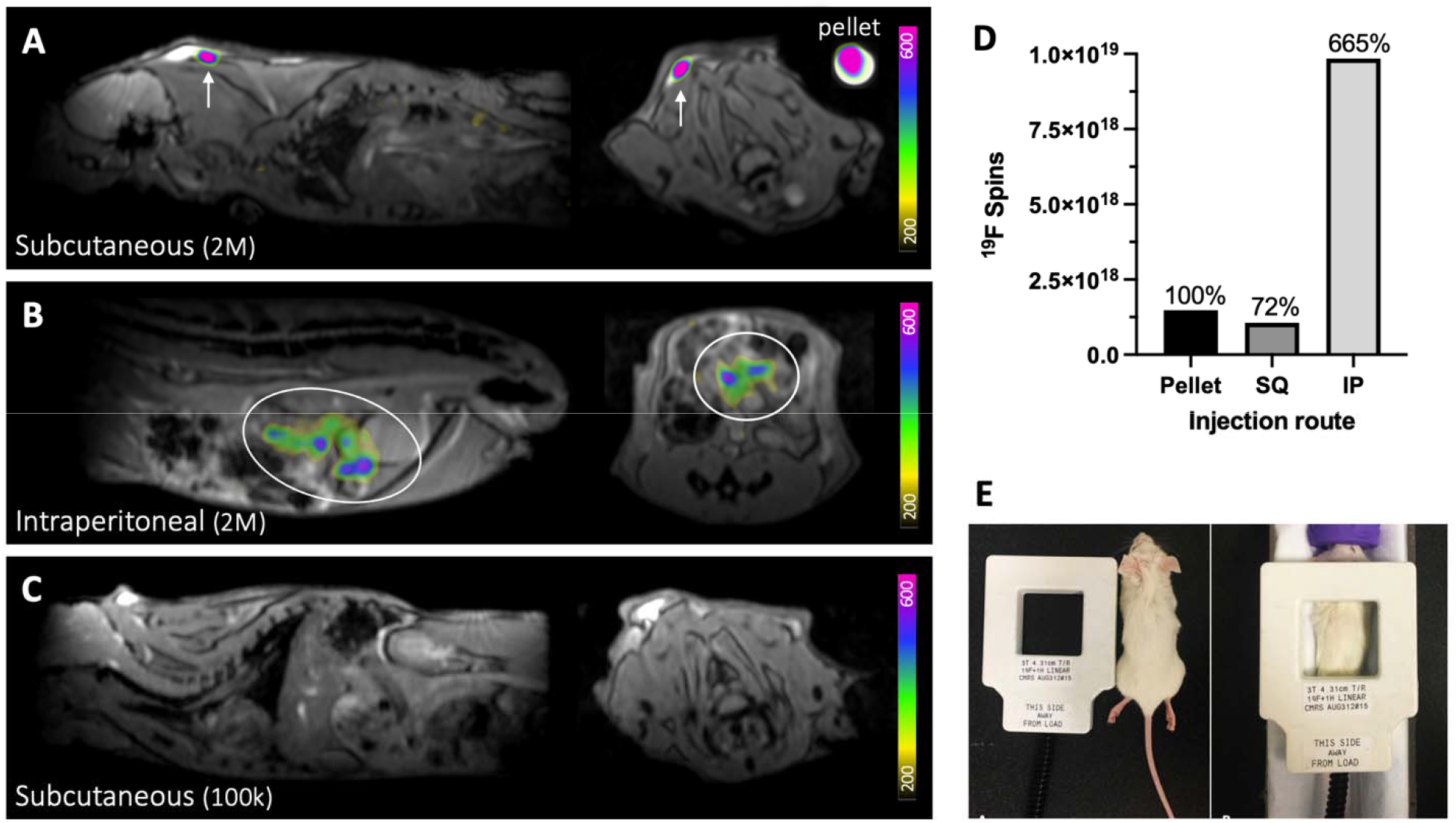
Detection of 2 × 10^6^ PFPE-labeled MSC *in vivo*. ^19^F signal is detected at the injection site following (**A**) subcutaneous injection (white arrows) and (**B**) intraperitoneal injection (white ovals). A cell pellet of the same cell number was imaged alongside the mouse for comparison (shown in **A**). (**C**) After subcutaneous administration, 1 × 10^5^ cells were undetected, as this cell number is below the detection threshold for MSC. Images are sagittal (left) and axial (right). M = million, k = thousand. (**D**) ^19^F signal measured from 2 × 10^6^ cells injected subcutaneous (SQ) was reduced compared to signal measured from the cell pellet, however, elevated ^19^F signal was measured from cells following intraperitoneal (IP) injection. (**E**) The dual-tuned surface coil is approximately the same size as a mouse and is placed directly over the injection site for imaging.

## Discussion

In this study, we began with an evaluation of *in vitro* sensitivity for MPI and ^19^F MRI of cells using ferucarbotran and PFPE nanoemulsions as labeling agents (respectively). Overall, fewer MSC were detected using MPI (4000) compared to ^19^F MRI (256000) using the same imaging time (1.5 minutes per cell pellet). Compared to ferucarbotran labeled MSC, more 4T1 cells were required for MPI detection (8000) as a result of lower cell uptake of ferucarbotran. These limits were defined with imaging criteria SNR > 5 and tested with 9 replicates to provide confidence.

These measurements of lowest cell number detected with MPI and ^19^F MRI are reasonable based on previous reports. As described earlier, Zheng et al. (2015)^5^ achieved detection of approximately 1000 ferucarbotran-labeled human embryonic stem cells (27 pg/cell, 27 *n*g) with SNR > 5. Our MSC detection limit was higher (4000 cells or 76 *n*g) and this is resulting from differences in MPI systems and acquisition. The amount of iron we detected (76 *n*g) is consistent with findings by Liu *et al.* (2021)^15^, where 64 *n*g ferucarbotran could be detected with mean SNR of 3.6, using another MOMENTUM MPI system and the same 2D imaging parameters. With longer imaging times in 3D, we demonstrated a minimum of 1000 ferucarbotran-labeled MSC (19 *n*g) could be visualized with SNR > 3. To the best of our knowledge, this is the lowest report of ferucarbotran detected from labeled cells. This study was the first to measure MPI detection limits for iron-loaded cancer cells (in 3D, as few as 4000 4T1 cells were detected, or 37 *n*g iron).

Previous investigation into ^19^F detection limits at 3T were conducted in dendritic cells (3.7 × 10^12^ ^19^F/cell)^7^ and macrophages (7.93 × 10^11^ ^19^F/cell)^9^, which have greater uptake of PFPE than what we measured in MSC (3.52 × 10^11^ ^19^F/cell). Ahrens *et al.* established an *in vitro* ^19^F MRI detection limit of 1.0 × 10^5^ dendritic cells/voxel on a 3T clinical scanner using a 2.5*SD_noise_ threshold^7^. Similarly, we achieved detection of 1.30 × 10^5^ MSC/voxel (1.5 minutes/pellet) and 8.55 × 10^4^ MSC/voxel in longer scans (4.5 minutes/pellet) using a 5*SD_noise_ threshold. In our previous work at 3 T, as few as 25,000 murine macrophages were detected (2.27 × 10^4^ cells/voxel) with longer imaging time^9^.

Cell labeling is fundamental in determining cellular sensitivity for both MPI and ^19^F MRI. We observed enhanced MPI detectability of MSCs compared to breast cancer cells, owing to increased endocytosis of SPION. Likewise, 4T1 cells did not label with PFPE sufficiently for ^19^F MRI detection. This result indicates that ^19^F may not be suitable for tracking labeled cancer cells to metastatic sites. Further improvements to PFPE nanoemulsions will enhance ^19^F cellular sensitivity, such as incorporation of paramagnetic agents^16,17^, or the addition of surface modifications to enhance the uptake of PFPE nanoemulsions^18^.

We recognize it is impossible to directly compare cell detection with MPI and ^19^F MRI due to their inherent differences, including structural configurations and imaging parameters. For this study, we attempted to optimize unique aspects of each modality in favor of sensitivity. MPI of cells was conducted using weak gradients (3 T/m) to increase the size of the FFR and enhance sensitivity. For FFL (line) MPI, the use of weak gradients (3 T/m) compared to stronger gradients (*e.g.* 6 T/m) expands the volume of FFR by 4 times. This leads to expected 4-fold enhancement in sensitivity, at the cost of reduced resolution (in this example, half resolution). It is also expected that signal averaging would improve sensitivity, however this has not been studied for MPI. In 2D, a significant reduction in background noise was measured with 8 averages compared to 1 average, however, this did not improve cell detection with MPI, as we defined it. 3D imaging using 35 projections did offer improvement in sensitivity for both ferucarbotran-labeled MSC (detection of 2000 cells) and breast cancer cells (4000 cells). Lastly, optimization of excitation amplitudes may lead to improved cell sensitivity; in this study we used 22 mT (X-channel) and 26 mT (Z-channel) by default.

^19^F MRI of PFPE-labeled cells was conducted at a clinical field strength (3 T). The implementation of the surface coil and optimized 3D bSSFP sequence is crucial to enable ^19^F cell tracking at 3 T. The theoretical optimal flip angle for ^19^F at 3 T is 72° and our investigation showed highest SNR was produced for flip angles between 60 - 80°. However, transmit/receive surface coils provide non-uniform sensitivity, due to spatial variations in applied energy and flip angle^19,20^. For this reason, cell pellets were imaged directly in the center of the coil to maximize sensitivity. Likewise, the surface coil was placed directly above the region of interest for *in vivo* ^19^F imaging. High signal averaging was used for ^19^F image acquisition (115) to improve SNR. Longer imaging times with 345 signal averages enabled detection of 3 additional pellets of 256 × 10^3^ cells and 3 of 9 pellets of 128 × 10^3^ PFPE-labeled MSC.

After assessing *in vitro* cell detection limits of MPI and ^19^F MRI, a preliminary assessment of *in vivo* detection factors was conducted. It has previously been shown that there is no attenuation of MPI signal from biological tissue^21,22^. In agreement, following subcutaneous injection of 2 × 10^6^ MSC, we measured only a small reduction in cell detection with MPI (5%). However, for a lower cell number (1 × 10^5^), there was reduction of signal measured *in vivo* compared to the pellet (by 47-85%, depending on injection route). Here we recognize that the dispersion of cells from the injection site reduces the cell density per voxel, leading some cells to fall below the intravoxel detection limit. MPI detection of MSC was most reduced after intravenous injection (85% reduction). It is expected that cells administered intravenously would be most disperse as they circulate through the venous circulation before accumulating in the lung capillaries (shown in **Figure 6A**). Similarly, previous work by Wang et al. (2020)^6^ showed that 1 × 10^5^ ferucarbotran-labeled stem cells could be detected with MPI *in vivo* following subcutaneous injection but not following intravenous injection. In our study, we could achieve detection of 1 × 10^5^ cells after intravenous injection, which can be attributed to the choice of gradient strength (3.0 T/m vs. 5.7 T/m).

Likewise for ^19^F MRI, dispersion of PFPE-labeled cells (1 × 10^6^) in patients was previously reported to limit the detectability of these cells following administration^7^. In our study, ^19^F signal detected from a cell pellet of 2 × 10^6^ PFPE-labeled MSC was higher than ^19^F signal from the same number of cells injected subcutaneously. Conversely, ^19^F signal measured from cells injected to the intraperitoneal space was overestimated. This result could be explained by the large quantification region, as ^19^F sensitivity per imaging voxel was 3.79 × 10^16^ ^19^F/mm^3^ for cells in a pellet, compared to 2.03 × 10^16^ ^19^F/mm^3^ for cells *in vivo*.

For both MPI and MRI, there are other important considerations which reduce detectability of cells *in vivo*. For MPI, this includes increased background signal associated with mouse digestion^23^. While we accounted for background signal in the mouse using signal subtraction, this technique is not permissible for longitudinal cell tracking studies. Background signal is variable across mice and day-to-day. In our experience, this can be reduced by mouse fasting, however, the amount of signal is unpredictable. Ultimately this background signal may obscure detection of cells, especially in low cell numbers^24^, as it is challenging (at this time, impossible) to distinguish the signal associated with cells from background. Second, there is some evidence that Brownian relaxation of SPION in different tissue environments may be altered, leading to reduced MPI sensitivity^25–27^. Brownian motion refers to the physical rotation of SPIONs; this motion is reduced in tissues with increased stiffness (e.g. muscle) which leads to increased Brownian relaxation times (thus, lower sensitivity and resolution). Ongoing work aims to determine whether this plays a role in the detection of SPION-labeled cells. Another consideration affecting ^19^F detection of PFPE-cells *in vivo* is the coil filling factor. The volume of a mouse is much larger than the volume of a cell pellet, thus SNR for detection of cells *in vivo* is expected to be reduced due to increased image noise. Lastly, mouse breathing motion can lead to blurring of signal which will reduce the maximum signal intensity associated with cells and could potentially render cells undetected by MRI and MPI.

## Conclusion

MPI has the potential to be more sensitive than ^19^F MRI for cell tracking. In this study, fewer MSC (4000 cells, 76 ng iron) were detected with MPI than ^19^F MRI (256000 cells, 9.01 × 10^16^ ^19^F atoms), using the same scan time. Furthermore, reduced ferucarbotran labeling was observed in 4T1 breast cancer cells compared to MSC, leading to a detection limit of 8000 breast cancer cells (74 ng iron). With longer imaging times, as few as 2000 MSC (38 ng ferucarbotran) and 4000 breast cancer cells (37 ng ferucarbotran) were detected with MPI and 128000 MSC (4.51 × 10^16^ ^19^F atoms) were detected with ^19^F MRI with SNR >5. Determination of these detection thresholds *in vitro* is useful to anticipate the minimum number of cells that are required for detection *in vivo*. However, we demonstrated that there are several factors *in vivo* which led to reduced detectability of cells, particularly the effect of cell dispersion which reduces cell density per imaging voxel. There is no doubt that cellular sensitivity for these modalities will continue to improve with further developments. It is essential to understand and improve cellular sensitivity to advance imaging of cellular therapeutics.

## Acknowledgements

We would like to acknowledge funding from the Natural Science and Engineering Council of Canada.

