## Supplementary Figures for "The sensitivity of magnetic particle imaging and fluorine-19 magnetic resonance imaging for cell tracking": Supplementary Figures.pdf

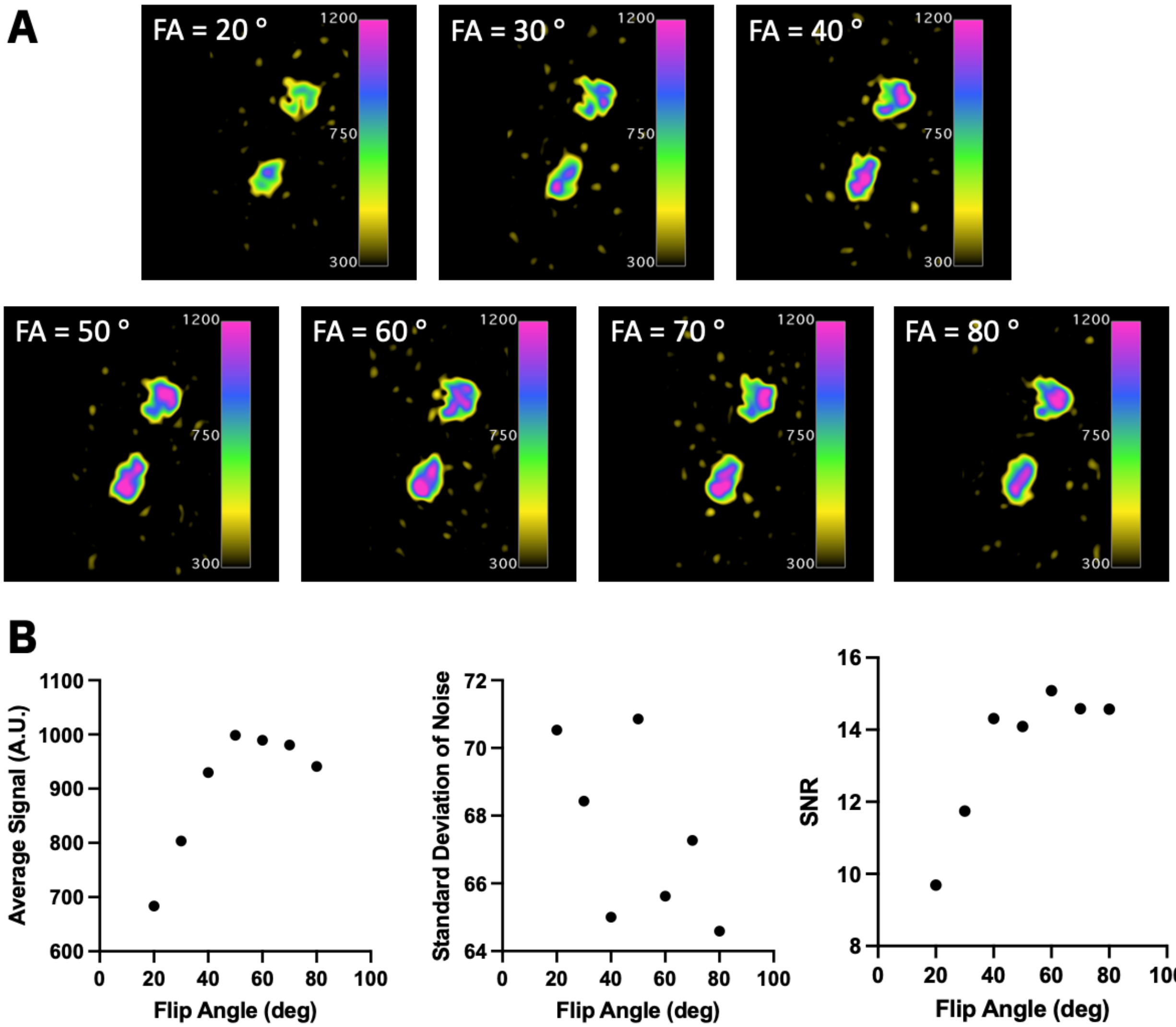


**Supplementary Figure 1. Optimization of $^{19}F$ flip angle at 3 T. (A)** Two reference phantoms (3.33 x $10^{16}$ $^{19}F$ spins/$\mu$L) were imaged using 3D bSSFP with 80 excitations and various flip angles: 20, 30, 40, 50, 60, 70, and 80 degrees. **(B)** The average signal of the reference tubes, the standard deviation of background noise, and the SNR were measured for each flip angle.

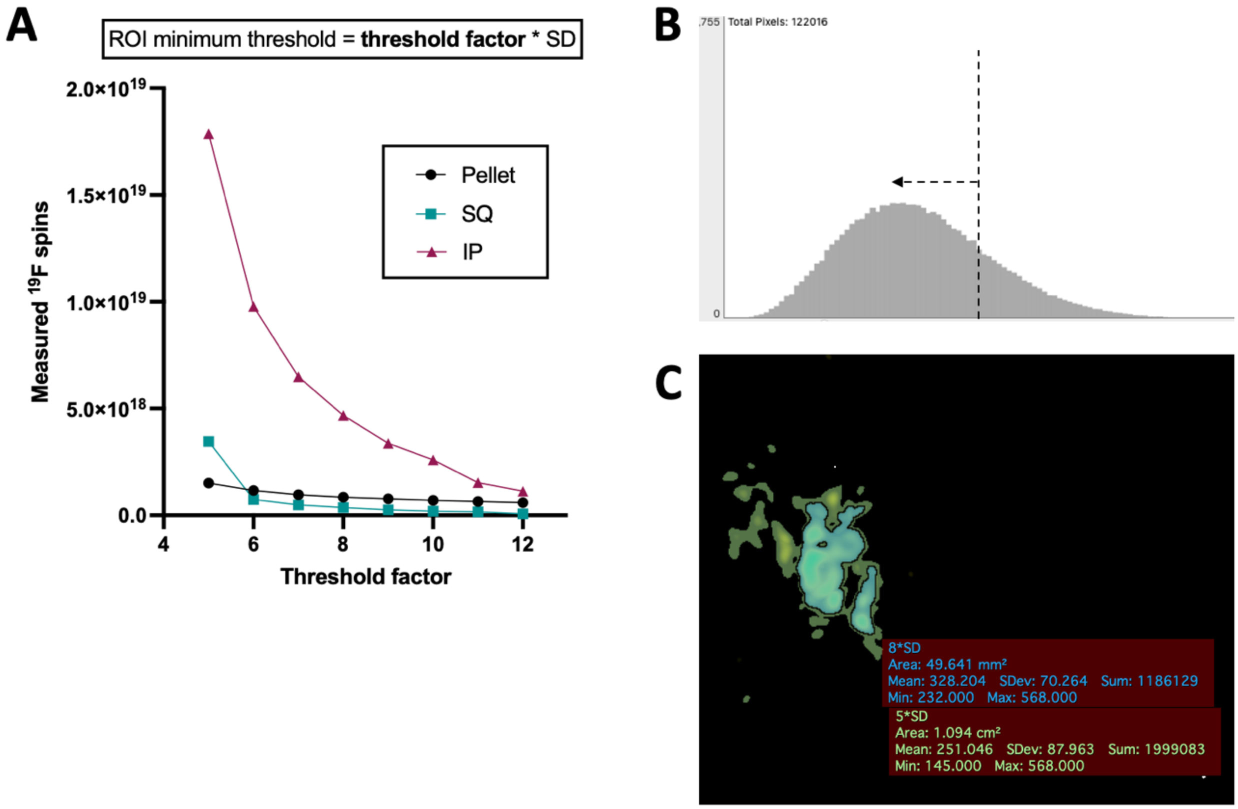


**Supplementary Figure 2. (A)** Various thresholds ($5*SD_{noise} – 12*SD_{noise}$) were used to segment $^{19}F$ signal from 2 x $10^6$ PFPE-labeled MSC, in a pellet or *in vivo* (subcutaneous, SQ; intraperitoneal; IP). These images are shown in Figure 7. Regardless of the threshold value used, $^{19}F$ signal measured from cells administered IP exceeded that of SQ and from pellets. In this quantification there is no correction applied to signal values. **(B)** Voxel distribution of background $^{19}F$ noise was non-Gaussian and left-skewed, indicating there is a Rician distribution of noise. **(C)** A demonstration of the segmentation method described in Bouchlaka et al. (2016)[14] from a single coronal slice of the mouse administered PFPE-cells IP (also shown in **Figure 7B**). $^{19}F$ signal between $5*SD_{noise}$ – $8*SD_{noise}$ (green) was corrected with a factor of 0.655 and $^{19}F$ signal exceeding $8*SD_{noise}$ (blue) was not corrected. Signal above $5*SD_{noise}$ was included for $^{19}F$ quantification.

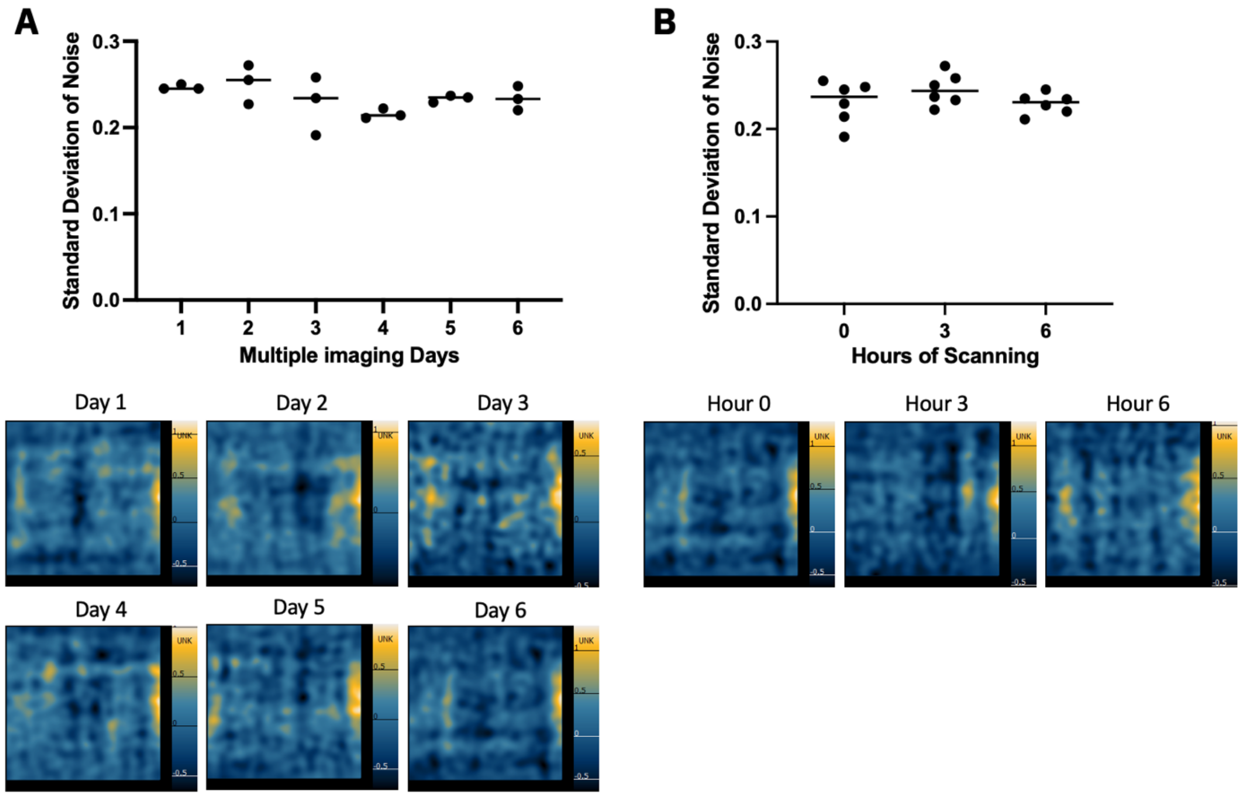


**Supplementary Figure 3.** The standard deviation of background noise ($SD_{noise}$) for MPI was measured from an empty sample holder at the beginning (t = 0h), middle (t = 3h), and end (t = 6h) of six imaging sessions. There were no significant differences in measured $SD_{noise}$ between **(A)** imaging days or **(B)** the 3 timepoints.
